# Bile-induced biofilm formation in *Bacteroides thetaiotaomicron* requires magnesium efflux by an RND pump

**DOI:** 10.1101/2023.07.26.550638

**Authors:** Anne-Aurélie Lopes, Julien Deschamps, Sonia Georgeault, Thomas Cokelaer, Romain Briandet, Jean-Marc Ghigo

## Abstract

*Bacteroides thetaiotaomicron* is a prominent member of the human gut microbiota contributing to nutrient exchange, gut function, and maturation of the host’s immune system. This obligate anaerobe symbiont can adopt a biofilm lifestyle and it was recently shown that *B. thetaiotaomicron* biofilm formation is promoted by the presence of bile, a process also requiring a *B. thetaiotaomicron* extracellular DNase, which is not, however, regulated by bile. Here we showed that bile induces the expression of several Resistance-Nodulation-Division (RND) efflux pumps and that inhibiting their activity with a global competitive efflux inhibitor impaired bile-dependent biofilm formation. We then showed that, among the bile-induced RND-efflux pumps, only the tripartite BT3337-BT3338-BT3339 pump, re-named BipABC (for Bile Induced Pump A (*BT3337*), B (*BT3338*) and C (*BT3339*), is required for biofilm formation. We demonstrated that BipABC is involved in the efflux of magnesium to the biofilm extracellular matrix, which leads to a decrease of eDNA concentration. The release of magnesium in the biofilm matrix also impacts biofilm structure, potentially by modifying the electrostatic repulsion forces within the matrix, reducing interbacterial distance and allowing bacteria to interact more closely and form denser biofilms. Our study therefore identifies a new molecular determinant of *B. thetaiotaomicron* biofilm formation in response to bile salts and provides a better understanding on how an intestinal chemical cue regulates biofilm formation in a major gut symbiont.

**IMPORTANCE:** *Bacteroides thetaiotaomicron* is a prominent member of the human gut microbiota able to degrade dietary and host polysaccharides, altogether contributing to nutrient exchange, gut function, and maturation of the host’s immune system. This obligate anaerobe symbiont can adopt a biofilm community lifestyle, providing protection against environmental factors that might, in turn, protect the host from dysbiosis and dysbiosis-related diseases. It was recently shown that *B. thetaiotaomicron* exposure to intestinal bile promotes biofilm formation. Here we reveal that a specific *B. thetaiotaomicron* membrane efflux pump is induced in response to bile, leading to the release of magnesium ions, potentially reducing electrostatic repulsion forces between components of the biofilm matrix. This leads to a reduction of interbacterial distance and strengthens the biofilm structure. Our study therefore provides a better understanding of how bile promotes biofilm formation in a major gut symbiont, potentially promoting microbiota resilience to stress and dysbiosis events.

## INTRODUCTION

*Bacteroides thetaiotaomicron* is a prominent human gut symbiont involved in beneficial nutrient exchanges, gut functions, and maturation of the host’s immune system from the earliest stages of life (1–5). *B. thetaiotaomicron* can degrade dietary and host polysaccharides and is often found adhering to food particles or grazing on the polysaccharide-rich gut mucus, developing biofilm-like communities that provide a stable and protective environment that also favors genetic exchange (2, 6–8). Several factors promoting the formation of *B. thetaiotaomicron* biofilm have been identified, including polysaccharide utilization receptors, capsular polysaccharides, and type V pili (9–12). We also recently showed that physiological concentrations of bile salts induce biofilm formation in a wide range of *B. thetaiotaomicron* strains and many Bacteroidales, suggesting that this relevant gut compound, beyond its critical digestive roles, could also induce biofilm formation *in vivo* (13, 14). Whereas bile-dependent biofilm formation requires the BT3563 DNase activity degrading extracellular DNA (eDNA) (14), the expression of *BT3563* gene is not induced by bile salts and the link between bile exposure and biofilm formation is therefore still unclear.

We and others previously showed that bile salts induce the expression of tripartite Resistance-Nodulation-Division (RND) efflux pumps in various bacteria, including *B. thetaiotaomicron* (14–18). Whereas this process could contribute to reduce bile toxicity, RND efflux pumps also actively transport a broad range of substrates (4, 17, 19–21). Here, we hypothesized that *B. thetaiotaomicron* RND-type efflux pumps could directly contribute to bile-dependent biofilm formation by secreting compounds into the biofilm matrix. We first showed that blocking RND-efflux pumps with the efflux inhibitor phenylalanine-arginine beta-naphthylamide (PAßN) reduced biofilm formation. By comparing gene expression in presence and absence of bile, we determined that bile induces the expression of seven *B. thetaiotaomicron* RND-efflux pumps and we showed that only the tripartite efflux pump encoded by the *BT3337*, *BT3338* and *BT3339* gene cluster is required for bile-dependent biofilm formation. We then demonstrated that this pump, renamed BipABC, exports magnesium divalent cations, which decreases the concentration of eDNA in the *B. thetaiotaomicron* extracellular matrix, allowing bacteria to interact more closely and form more compact biofilm structures. The demonstration of a direct link between exposure to bile and the expression of a specific RND-type pump and magnesium efflux provides new insights into the regulation of biofilm formation in *B. thetaiotaomicron*.

## RESULTS

### Inhibiting *B. thetaiotaomicron* RND efflux pumps impairs bile-dependent biofilm formation

To evaluate the contribution of RND-efflux pumps to *B. thetaiotaomicron* VPI-5482 biofilm formation in presence of bile salts, we tested the impact of blocking its RND-type efflux pumps with the competitive efflux pump inhibitor phenylalanine-arginine ß-naphthylamide (PAßN) (22, 23). We observed that the addition of 25 µg/mL PAßN significantly inhibited bile-dependent biofilm formation (Fig. 1A), while addition of PAßN did not affect growth or viability (Supplementary Fig. S1AB). We also verified that PAßN did not increase membrane permeability, since exposure to PAßN did not modify *B. thetaiotaomicron* VPI-5482 sensitivity to vancomycin, an antibiotic unable to cross the outer membrane of Gram-negative bacteria unless permeabilized (Supplementary Table S1) (24). Finally, we used Hoechst H33342 as a probe for efflux activity and a real-time drug accumulation assay (25) to demonstrate the specific impact of PAßN on efflux. We first showed that, compared to the PBS control, the presence of bile decreased the intracellular concentration of Hoechst H33342, indicative of its increased efflux (Fig. 1B). By contrast, dual exposure to bile and PAßN increased the intracellular accumulation of Hoechst H33342, indicating that PAßN blocked bile-induced RND-type efflux (Fig. 1B). Taken together, these results indicated that *B. thetaiotaomicron* VPI-5482 RND-type efflux contributes to bile-dependent biofilm formation.

**Figure 1:**
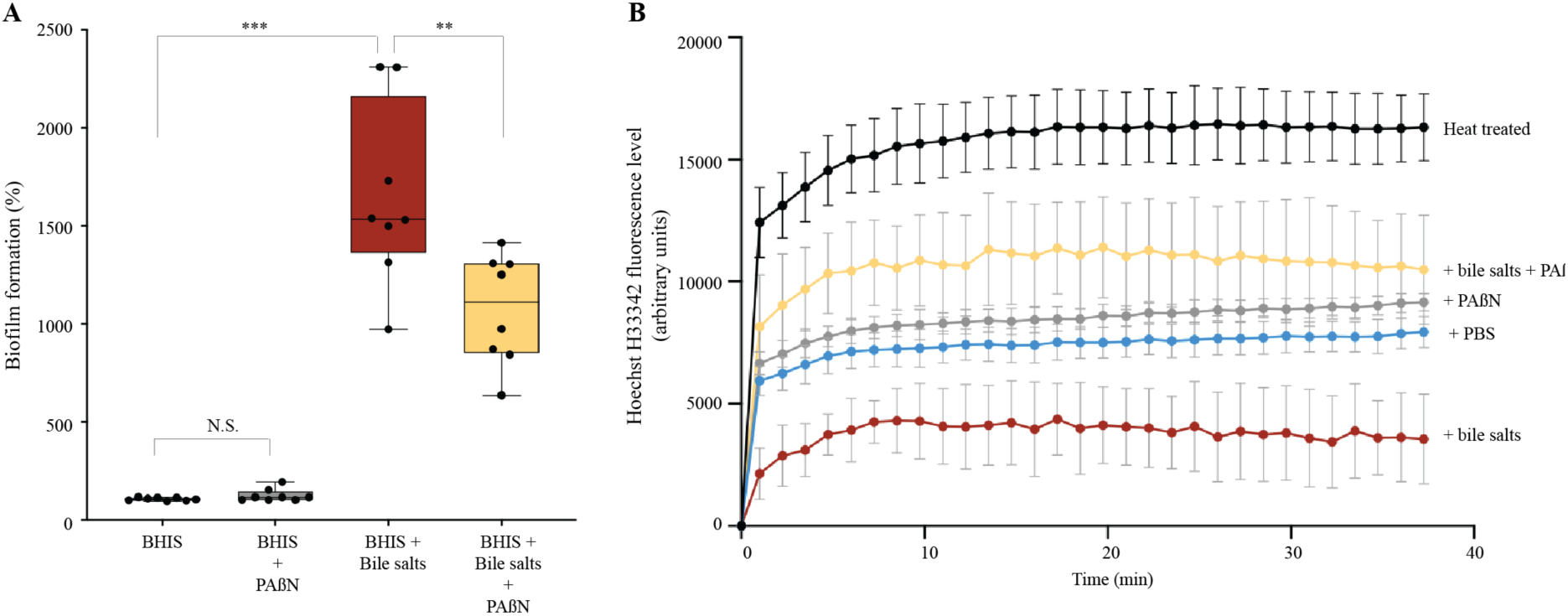
RND-type efflux is involved in bile-dependent biofilm formation. A. 96-well plate crystal violet assay of *B. thetaiotaomicron* VPI-5482 biofilm formation after 48h growth in BHIS in the absence and presence of phenylalanine-arginine ß-naphthylamide (PAßN) (25 µg/mL) and/or 0.5% bile salts (BS). Mean of WT biofilm formation in BHIS was adjusted to 100%. Min-max boxplot of 8 biological replicates, each of them being the mean of three technical replicates, for each condition. NS: non-significant. ** p-value<0.005; *** p-value<0.0005. Statistics correspond to an unpaired, nonparametric Mann–Whitney *U* test. **B.** Hoechst H33342 accumulation assay (2.5 µM) in *B. thetaiotaomicron* VPI-5482 in the absence and presence of 0.5% bile salts (BS) and/or PAßN (25 µg/mL). Hoechst H33342 level in heat-inactivated bacteria represent the maximum fluorescence level. Mean of 4 biological replicates.

### The RND efflux pump BT3337-3339 is required for bile-dependent biofilm formation

To identify which specific *B. thetaiotaomicron* efflux pumps could be involved in biofilm formation, we performed a RNAseq analysis to compare the expression of the 21 gene clusters annotated as RND-type efflux pumps in the VPI-5482 genome in absence or presence of bile (26). This analysis confirmed the previously observed induction of the tripartite BT2793-2795 efflux pump (14), along with 6 other RND efflux pumps clusters, which were induced close to or above a 2-fold factor in bile salt conditions (namely *BT0297-0300, BT1965-1967, BT2117- 2119, BT2686-2688, BT2835* and *BT3337-3339*) (Supplementary Table S2 and S3). We then deleted each of these 7 RND-type efflux pump operons and showed that only the deletion of the *BT3337-3339* gene cluster led to a reduction of biofilm capacity to a level similar to the one obtained upon the addition of PAβN in the wild-type (WT) strain. Moreover, the addition of PAβN did not further reduce biofilm formation of the Δ*BT3337-3339* mutant (Fig. 2). All other RND-type efflux pump mutants either only mildly decreased or even increased (*BT1965-67*) biofilm formation (Fig. 2). These results indicated that the involvement of the BT3337-3339 RND-type efflux pump in biofilm formation in presence of bile and this pump was hereafter renamed BipABC, for Bile-Induced Pump A (*BT3337*), B (*BT3338*) and C (*BT3339*).

**Figure 2:**
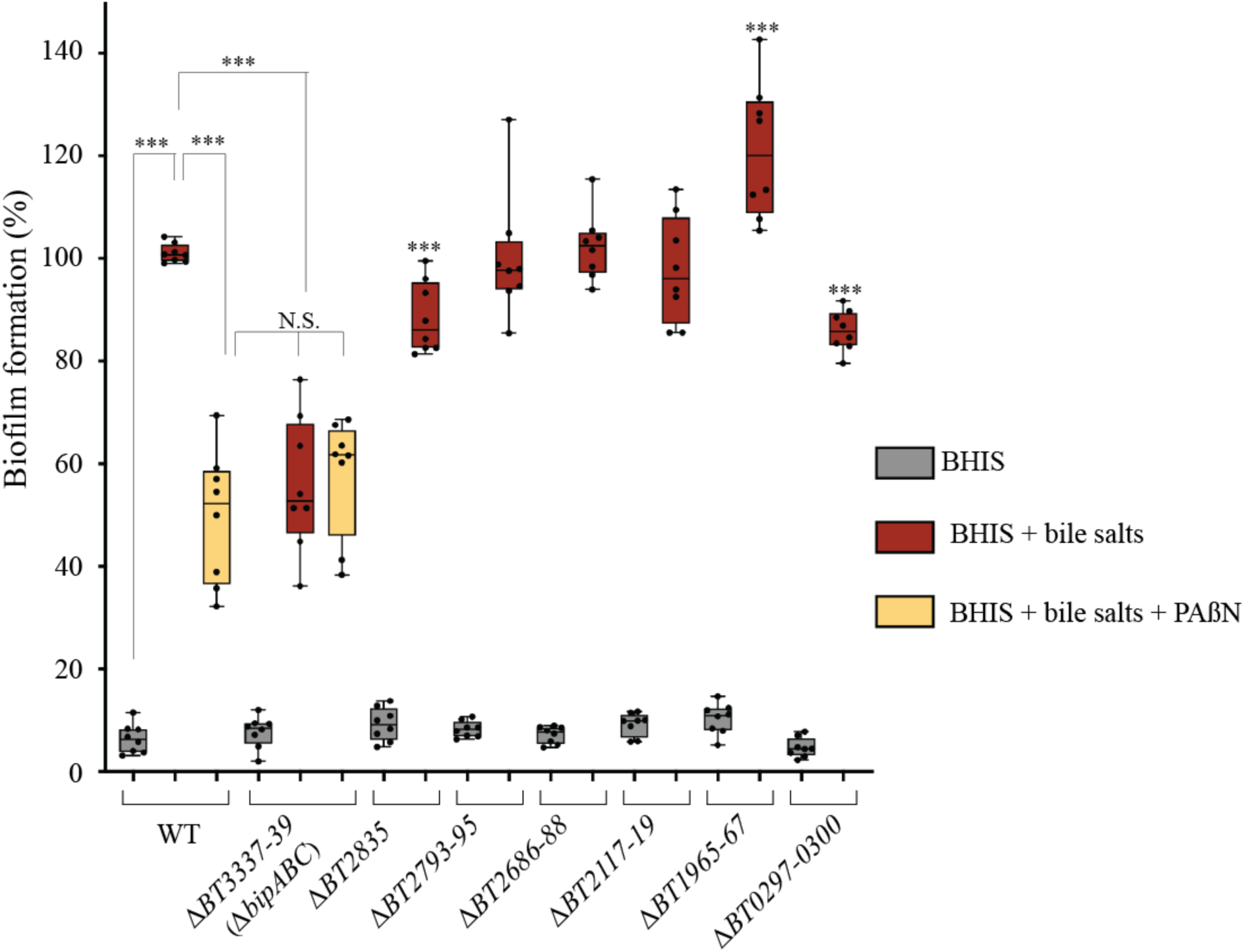
The deletion of *BT3337-39* gene cluster reduced *B. thetaiotaomicron* VPI-5482 biofilm formation. 96-well plate crystal violet biofilm assay after 48h growth in BHIS in the absence and presence of 0.5% bile salts (BS) of strains in which each of the 7 targeted RND- type efflux pumps clusters were deleted. VPI-5482 wild type (WT) and *ΔBT3337-39* biofilm formations were also compared without or with 25µg/mL of PAßN. Mean of WT in BHIS with 0.5% BS was adjusted to 100%. Min-max boxplot of 8 biological replicates, each of them being the mean of six technical replicates, for each condition. *** p-value<0.0005; NS: non- significant compared to WT in BHIS with BS. Statistics correspond to an unpaired, nonparametric Mann–Whitney *U* test.

### The *B. thetaiotaomicron* eDNA content increases in absence of the BipABC efflux pump

To investigate how *B. thetaiotaomicron* efflux pumps contributed to bile-dependent biofilm formation, we compared the amount of extracellular DNA (eDNA), proteins and polysaccharides in the extracellular matrix of biofilms formed in absence and presence of PAßN. After normalization to the extent of recovered biomass, we observed that the addition of bile salts increased the concentrations of all three matrix components (Figure 3A and Supplementary Figure S2). By contrast, compared to the addition of bile salts alone, the addition of bile salts and PAßN led to an increase in eDNA, which was also observed in matrix extracted from a *ΔbipABC* mutant (Figure 3A). To test whether this eDNA increase could result from a reduction of eDNA degradation, we exposed for 24h the *B. thetaiotaomicron* genomic DNA to bacterial-free filtered culture supernatants of VPI-5482 WT and its *ΔbipABC* mutant with bile or bile and PAßN. As a control, we exposed genomic DNA to filtered culture supernatants of a Δ*BT3563* mutant lacking the previously described BT3563 extracellular DNase (14), which did not lead to eDNA degradation (Fig. 3B). All other conditions showed WT level of *B. thetaiotaomicron* genomic DNA degradation (Fig. 3B) suggesting that the lack of the BipABC efflux pump did not reduce the level of DNase activity in the tested supernatants (Fig. 3B).

**Figure 3:**
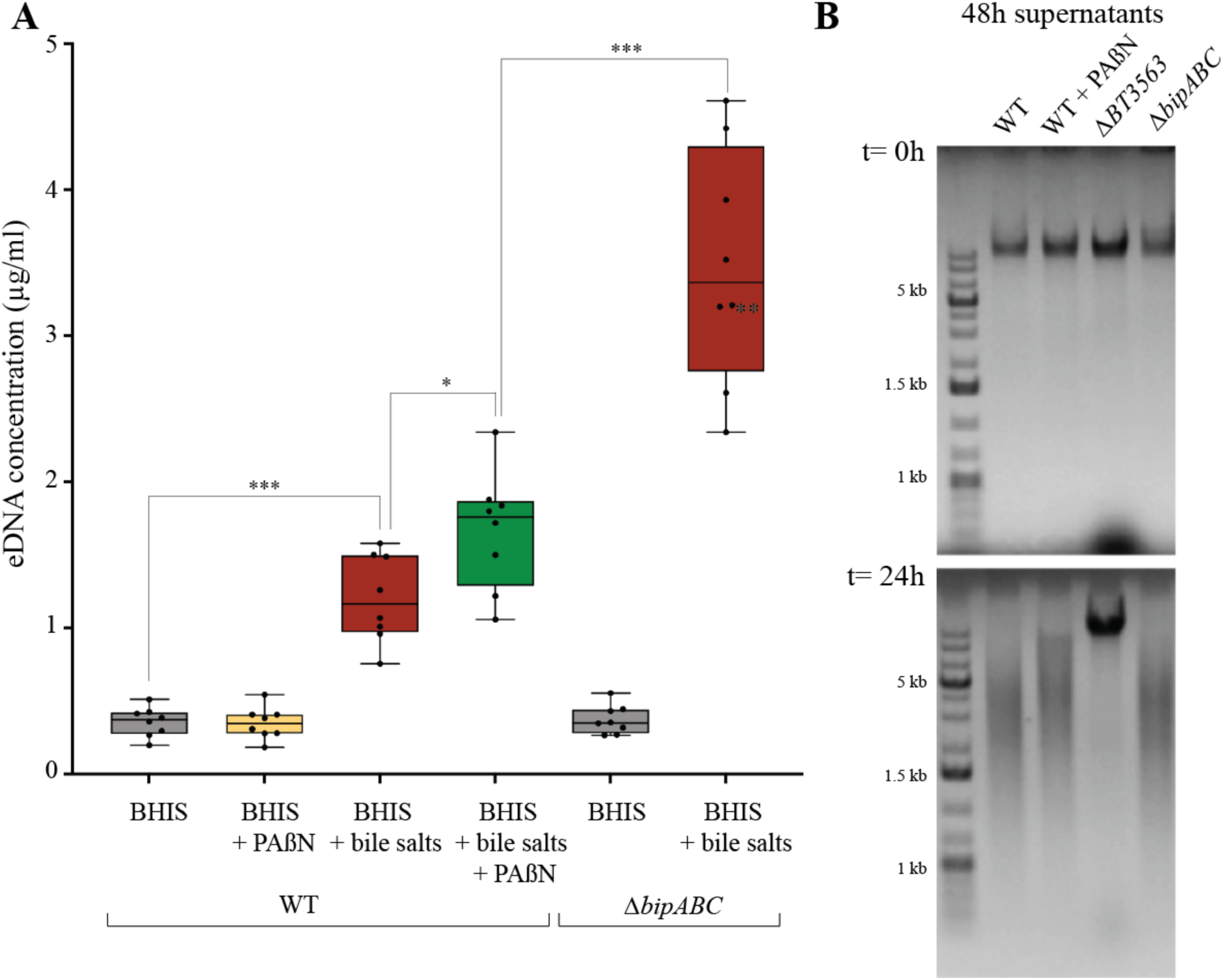
Addition of PAßN and deletion of *bipABC* (*BT3337-39)* genes lead to eDNA increase without impairing the BT3563 extracellular DNase activity. A. Concentration of DNA in the extracellular matrix of *B. thetaiotaomicron* VPI-5482 WT or its corresponding Δ*bipABC* mutant with 0.5% bile salts (BS) and 25 µg/mL of PAßN as indicated. Min-max boxplot of 8 biological replicates for each condition. * p-value<0.05; *** p-value<0.0005. Statistics correspond to an unpaired, nonparametric Mann–Whitney *U* test. **B.** Incubation of *B. thetaiotaomicron* genomic DNA (800 ng/mL final concentration) with the supernatant of 48h bile-free indicated cultures, and loaded immediately on a 1% agarose gel (t= 0 h) and after 24h incubation (t= 24 h) at 37 °C.

### Addition of magnesium restores *B. thetaiotaomicron* bile-dependent biofilm in the Δ*bipABC* efflux pump mutant

Among potential substrates exported by RND-type efflux pumps, divalent cations have been reported to impact the interactions between biofilm matrix components (27, 28). To test whether impairing cation efflux could account for the biofilm defect observed in the Δ*bipABC* efflux pump mutant, we first evaluated the impact of EDTA, a chelator of a wide range of divalent cations, on bile-dependent biofilm formation. We showed that extracellular addition of EDTA inhibited bile-dependent biofilm formation reaching a level similar to PAßN inhibition at the non-toxic EDTA concentration of 0.2mM (Fig.4A and Supplementary Figure S3A). Among the cations potentially chelated by EDTA, we showed that addition of non-toxic concentration of magnesium (35-50 mM) restored wild-type level of biofilm formation in presence of PAßN (Fig. 4B and Supplementary Figure S3B). Consistently, addition of magnesium also restored biofilm formation of the *ΔbipABC* mutant (Fig. 4C and Supplementary Figure S3C). By contrast such a chemical complementation was not observed with addition of calcium, cadmium, cobalt, iron, kalium or zinc (Supplementary Figure S4 A-F). Finally, we also confirmed that magnesium concentration was increased in the extracellular matrix of the WT strain *B. thetaiotaomicron* VPI-5482 grown in presence of bile salts but decreased, compared to WT and bile salts, when grown in presence of bile salts and PAßN (Fig. 4D) as well as in the extracellular matrix of the *ΔbipABC* mutant in the presence of bile salts (Fig. 4D). These results further demonstrated that genetic inactivation or chemical inhibition of BipABC reduces magnesium-efflux and affects bile-dependent biofilm formation.

**Figure 4:**
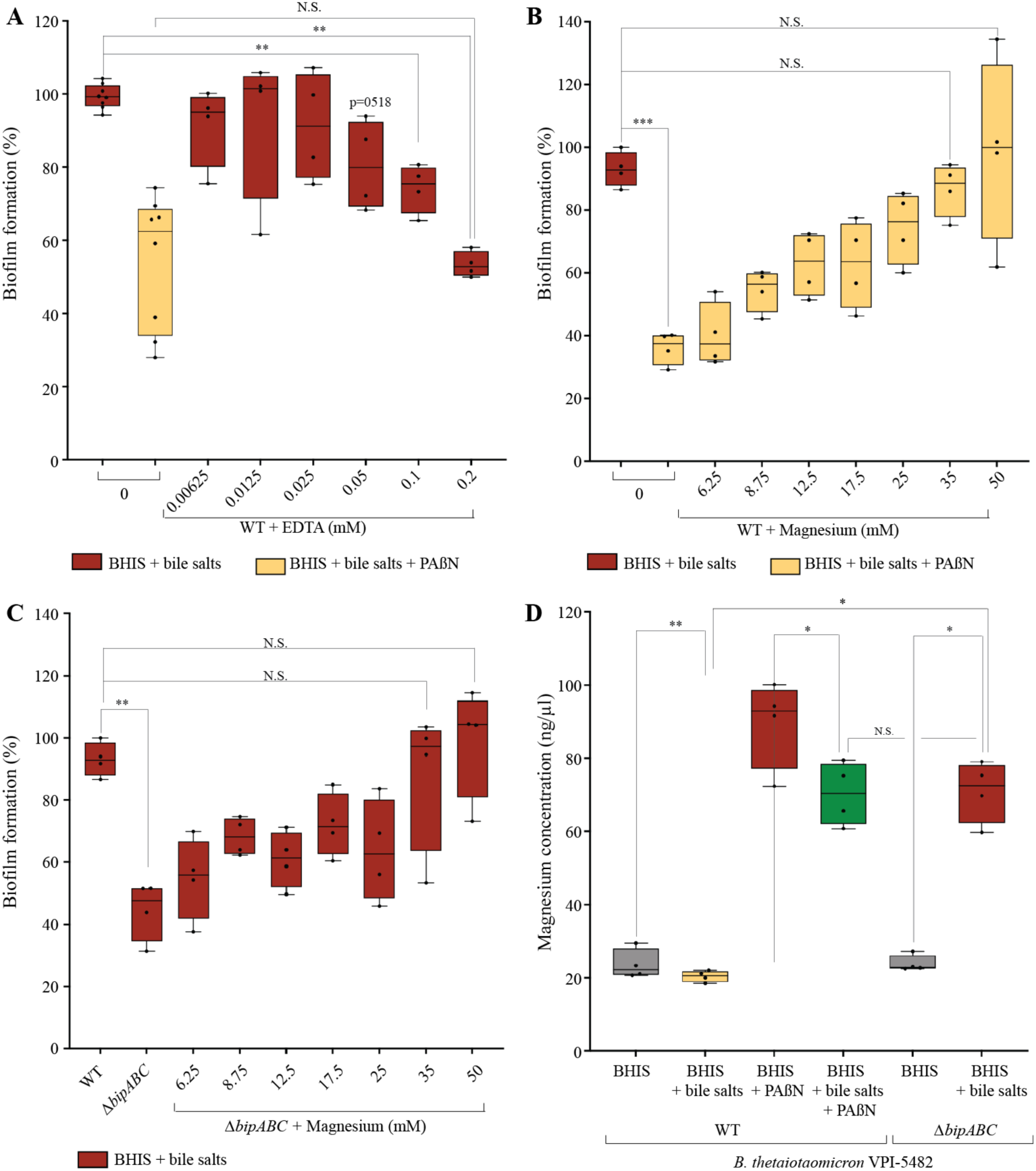
Addition of magnesium divalent ions restore RND efflux pump defect in bile-dependent biofilm formation in *B. thetaiotaomicron*. A and **B**96-well plate crystal violet biofilm assay after 48h growth in BHIS of VPI-5482 in the presence of 0.5% bile salts without or with 25µg/mL of PAßN and different non-toxic concentrations of EDTA (**A**) or Magnesium (**B**). Mean of WT in BHIS with 0.5% BS was adjusted to 100%. Min-max boxplot of 8 biological replicates for each condition. **C.** 96-well plate crystal violet biofilm assay after 48h growth in BHIS of Δ*bipABC* in the presence of 0.5% bile salts and different non-toxic concentrations of magnesium. Mean of WT in BHIS with 0.5% BS was adjusted to 100%. Min- max boxplot of 8 biological replicates for each condition. **D.** Quantification of magnesium concentration in the ECM in VPI-5482 without and with 0.5% bile salts or 25 µg/mL of PAßN and in Δ*bipABC* mutants without and with 0.5% BS. Min-max boxplot of 8 biological replicates for each condition. * p-value<0.05; ** p-value<0.005; *** p-value<0.0005; N.S.: non- significant. Statistics correspond to an unpaired, nonparametric Mann–Whitney *U* test.

### Magnesium efflux decreases the concentration of extracellular DNA in *B. thetaiotaomicron* biofilm

In addition to restoring bile-dependent biofilm formation, we showed that extracellular addition of magnesium also reduced the concentration of eDNA in the extracellular matrix of WT *B. thetaiotaomicron* VPI-5482 grown in presence of bile salts and PAßN as well as a in *ΔbipABC* mutant grown in presence of bile salts (Fig. 5A). To investigate the effect of magnesium on eDNA, we use electrophoresis profiling to compare the size of eDNA fragments extracted from the extracellular matrix of biofilms formed by WT VPI-5482 and the Δ*bipABC* and Δ*BT3563* DNase mutant in presence of bile salts with and without PAßN and magnesium. This analysis revealed that a high concentration of DNA fragments sized above 600 bp in the WT strains as well as in the mutants grown in presence of bile (Fig. 5B). By contrast, addition of magnesium eliminated this fraction above 600 bp eDNA fraction (Fig. 5B), suggesting that magnesium divalent cations could either activate eDNA degradation by other *B. thetaiotaomicron* DNases than BT3563 or limit the release of eDNA in the biofilm matrix.

**Figure 5:**
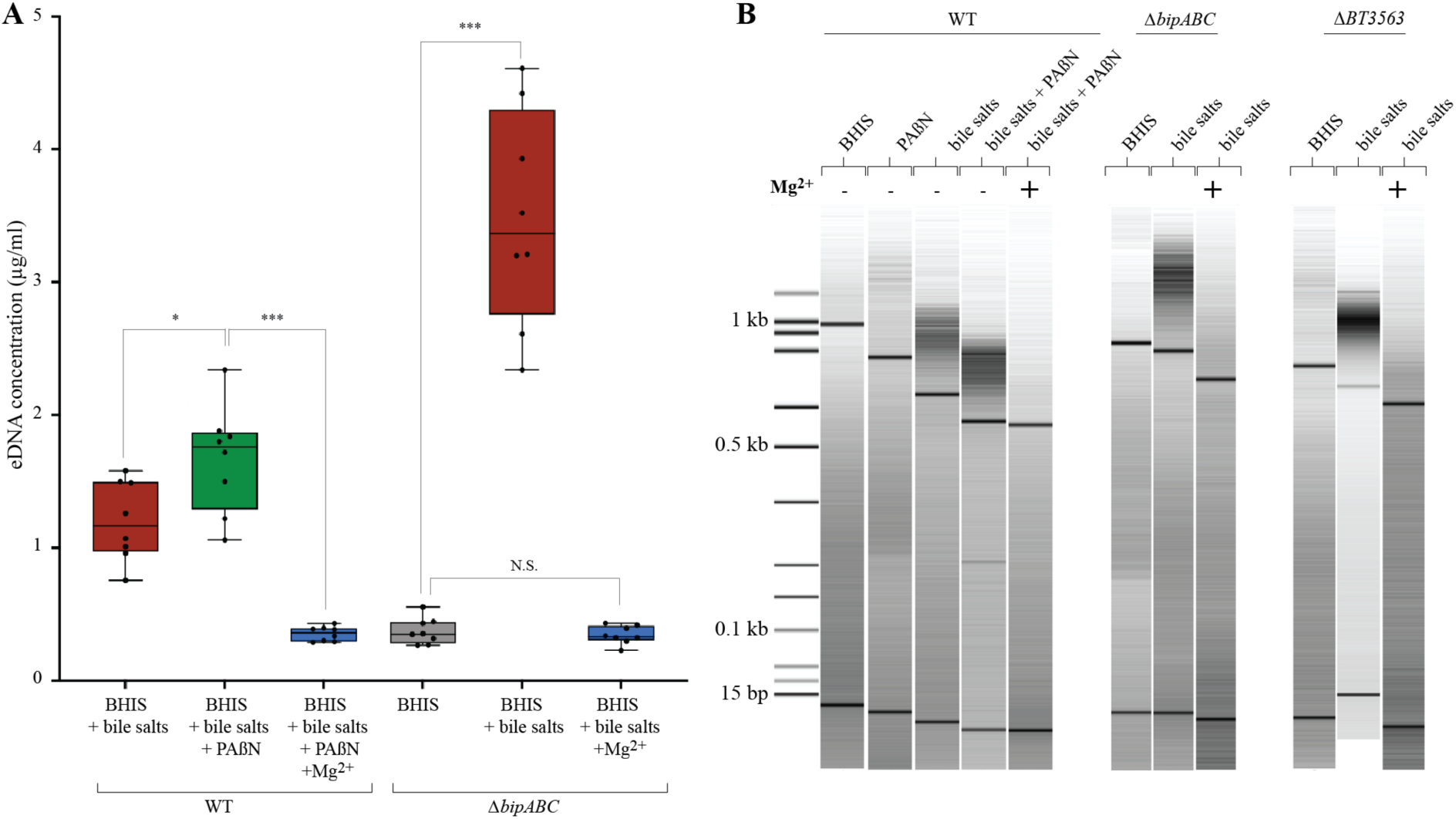
Quantification and effects of magnesium on eDNA concentration and profile. **A.** Concentration of DNA in the extracellular matrix of VPI-5482 without and with 0.5% bile salts or 25 µg/mL of PAßN or 35mM magnesium and in deleted mutants without and with 0.5% bile or 35mM magnesium. Min-max boxplot of 8 biological replicates for each condition. * p- value<0.05, *** p-value<0.0005. N.S.: non-significant. Statistics correspond to an unpaired, nonparametric Mann–Whitney *U* test. **B.** Bioanalyzer DNA profile of extracellular matrix extract samples.

### Magnesium efflux modifies biofilm structure by reducing interbacterial distance

To test the impact of magnesium efflux on the structure of *B. thetaiotaomicron* biofilm, we used transmission electron microscopy (TEM) performed on ultra-thin section of resin-embedded biofilms (29). We observed that, compared to WT, the inhibition of efflux with PAßN or the deletion of the Δ*bipABC* efflux pump genes led to a significant decrease of biofilm bacterial density correlating with an increased interbacterial distance (Fig. 6). This phenotype could be complemented by the addition of magnesium to VPI-5482 in presence of PAßN or to the Δ*bipABC* mutant (Fig. 6). These results are consistent with the determination of biofilm biovolume after imaging of 48-h 96-well plate biofilms by confocal laser scanning microscopy, showing that blocking all RND efflux pump (PAßN condition), or only BipABC in presence of bile increased biovolume, which was corrected upon addition of magnesium (Supplementary figure S5). These results demonstrate the key role played by magnesium efflux and the bile- dependent BipABC efflux pump in the formation of dense biofilm in presence of bile.

**Figure 6:**
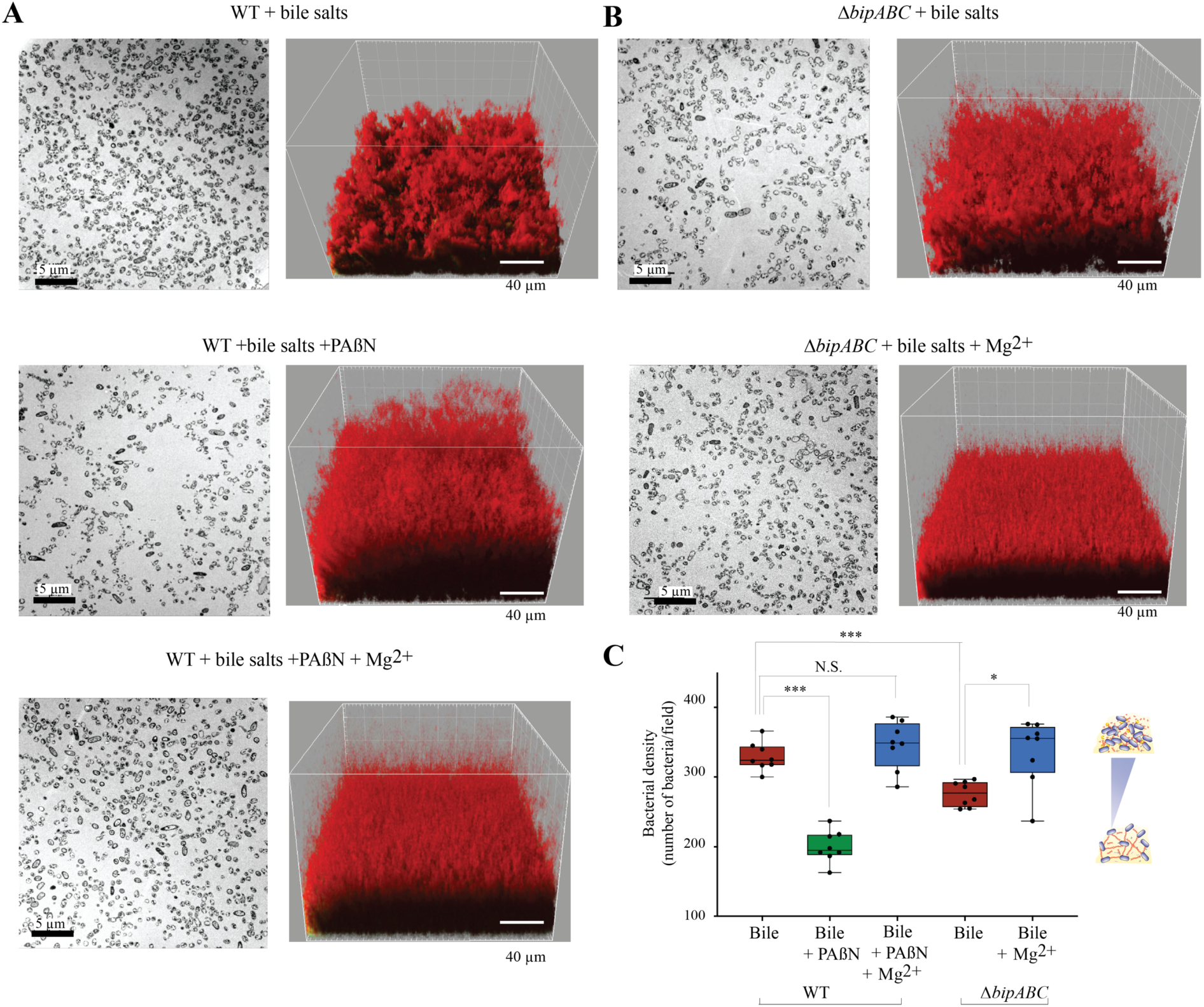
Increased interbacterial distance upon addition of PAßN and deletion of the BipABC RND-type pump is complemented by the addition of magnesium. A. Visualization of biofilms from *B. thetaiotaomicron* VPI-5482 WT and **B.** Visualization of biofilms from the Δ*bipABC* deletion mutant. **Left:** Transmission electron microscopy (TEM) images of ultra-thin sections of resin-embedded biofilms (scale bar: 5µm). **Right**: Imaris three-dimensional (3D) reconstruction of the biofilm from Confocal Laser Scanning Microscopy (CLSM) (scale bar: 40 µm). Bacterial cells are labeled in red with 5µM of SYTO61, a cell permeant nucleic acids dye. Culture conditions correspond to 0.5% bile salts, 25 µg/mL of PAßN or 35mM magnesium **C**. Interbacterial distance evaluated by the number of bacteria by observation field of ultra-thin section of TEM. Min-max boxplot of 8 biological replicates for each condition. * p-value<0.05; *** p-value<0.0005; N.S.: non-significant. Statistics correspond to an unpaired, nonparametric Mann–Whitney *U* test.

## DISCUSSION

In this study, we showed that bile salts induce the expression of several *B. thetaiotaomicron* RND-efflux pump, known to be involved in the efflux of a wide range of substrates (17, 21, 30). We then demonstrated that only the deletion of one bile-induced pump composed of BT3337, BT3338 and BT3339 – renamed BipABC – decreased biofilm formation and is therefore a new determinant of *B. thetaiotaomicron* biofilm formation.

In addition to BipABC, we showed that six other *B. thetaiotaomicron* RND-efflux pumps were up-regulated in presence of bile, however, none of them played a role in biofilm formation, suggesting that these pumps could mediate other adaptative phenotypes. RND-efflux pumps are tripartite protein complexes composed of an inner pump, determining the substrate specificity, a periplasmic adapter, and a passive outer membrane channel (17). These pumps mediate the efflux of various metabolites, and some of the identified pumps could for instance mediate the efflux of bile out of the cell as a tolerance mechanism against bile cytotoxic and detergent properties, as already demonstrated for *BT2793-2795* pump in *B. thetaiotaomicron* (31) and for RND-type efflux pumps in several enteric bacteria (18, 32, 33).

Besides bile-induced stress tolerance, the observed gene expression changes in presence of bile suggests that even at non-inhibitory concentration, bile is sensed by *B. thetaiotaomicron* and can deeply remodel its physiology. Induction of biofilm formation by bile could for instance be used as a host-derived signal to switch from planktonic to a biofilm lifestyle to stably colonize the gut or optimized food foraging in nutrient and bile-rich region of the digestive tract. We do not yet know how is bile sensed by the cell, nor how it translates into mediating efflux pump expression. Interestingly, we showed that bile also modulates the transcription of some AraC-family transcriptional regulators, which are known regulators of RND efflux pumps, with both positive and negative impact on biofilm formation (15, 18, 34, 35) and we are currently investigating their potential contribution to *B. thetaiotaomicron* biofilm formation.

By contrast with the known scaffolding role of eDNA in many bacterial species (14, 27, 28, 36), *B. thetaiotaomicron* biofilm formation in presence of bile was previously shown to involve the degradation of extracellular DNA (eDNA) by the DNase BT3563 released in the extracellular matrix (14). Here, we demonstrate that addition of magnesium to the extracellular matrix, either through BipABC pump activity or through direct magnesium supplementation, led to lower eDNA concentration in the biofilm matrix and increased biofilm formation, confirming that excess of eDNA impairs *B. thetaiotaomicron* biofilm formation. Since many DNases require divalent cation for activity (37), we hypothesized that BipABC-dependent magnesium efflux could increase the eDNA degradation by DNases. However, we showed that extracellular DNase activity on *B. thetaiotaomicron* genomic DNA was not impaired in a *bipABC* mutant, or in presence of the pump inhibitor PAβN. Moreover, our analysis of eDNA profile showed that, even in a *ΔBT3563* mutant, addition of magnesium leads to the disappearance of the above 600 bp DNA fraction. This suggests that magnesium could reduce eDNA concentration through an unidentified DNase whose activity in not detectable in our general extracellular DNase assay (Figure 3B). Alternatively, magnesium efflux in the biofilm matrix could limit the release of eDNA rather than impacting eDNA degradation.

One of the contributions of eDNA to biofilm architecture lies on its capacity to interact with charged components of the biofilm matrix (27, 38–40). We previously hypothesized that eDNA degradation by BT3563 could prevent or reduce electrostatic repulsion between an interbacterial grid of negatively charged eDNA molecule and negatively charged *B. thetaiotaomicron* surface and lead to denser biofilm structure (14). The high eDNA concentration associated to blocking magnesium efflux bloc would therefore increase the interbacterial distance between cells, whereas addition of extracellular magnesium restored WT distance between bacteria and strengthened biofilm formation. However, it is yet unclear whether magnesium efflux only reduces eDNA concentration in the matrix, allowing the formation of denser biofilm, or whether it directly interacts with cells and/or eDNA in the matrix and mediates electrostatic repulsion regardless of its impact on eDNA concentration. Indeed, magnesium divalent cations are important electrostatic component in biofilms (41, 42) and the requirement for its BipABC-dependent efflux suggests that, in addition to, or independently of eDNA degradation, efflux of magnesium could neutralize the negative charge conferred by degraded or undegraded eDNA phosphate-backbone groups (27, 28). By reducing the electrostatic repulsion forces between the matrix components, including those exposed before and after eDNA degradation, this could change the biofilm viscoelastic properties and allow closer interactions between bacteria, forming thicker and more compact biofilm structures (27).

In conclusion, our study provide evidence for a direct link between exposure to bile salts addition and factors necessary for *B. thetaiotaomicron* biofilm formation. Since biofilm formation likely provides a protective environment against environmental factors, the identification of BipABC-mediated magnesium efflux as a mechanistic link between an intestinal cue such as bile and *B. thetaiotaomicron* biofilm could lead to original strategies to foster biofilm formation by this important gut symbiont and potentially promote microbiota resilience to stress and dysbiosis events.

## MATERIALS AND METHODS

### Bacterial strains and growth conditions

*B. thetaiotaomicron* strains were grown in BHIS broth (43) supplemented with bile extract from bovine and ovine (Sigma, B8381) at 0.5%, phenylalanine-arginine ß-naphthylamide (PAßN) at 25µg/mL, magnesium at 35mM, unless indicated otherwise, EDTA, calcium, iron, kalium, cobalt, cadmium or zinc. Cultures were incubated at 37°C in anaerobic conditions in a C400M Ruskinn anaerobic-microaerophilic station. All media and chemicals were purchased from Sigma-Aldrich unless indicated otherwise. All experiments and genetic constructions of *B. thetaiotaomicron* were made in VPI-5482Δtdk background (44), which was developed for a two-step selection procedure of unmarked gene deletion by allelic exchange and has become the reference strain used in most studies. Therefore, VPI-5482Δtdk is referred to as wild type or VPI-5482 in this study.

### Construction of *B. thetaiotaomicron* mutants

For the generation of deletion strains by allelic exchange in *B. thetaiotaomicron*, we used the pLGB13 vector (45). Briefly, 1-kb upstream and downstream regions of the target sequence were cloned into the pLGB13 vector and transformed into the *E. coli* S17 λpir strain, which was used to deliver the vector to *B. thetaiotaomicron* by conjugation. Conjugation was carried out by mixing exponentially grown cultures of the donor and the recipient strains at a 2:1 ratio and placing the mixture on BHIS agar plates at 37°C under aerobic conditions overnight. After this, bacteria were plated on selective BHIS agar supplemented with erythromycin for selection of *B. thetaiotaomicron* transconjugants that underwent the first recombination event and gentamicin to ensure the exclusion of any *E. coli* growth. Erythromycin- and gentamicin-resistant colonies of *B. thetaiotaomicron* were then subjected to a second round of selection on BHIS agar plates supplemented with anhydrotetracycline (aTC) for selection of double recombined colonies. The resulting deletion mutants were confirmed by PCR and sanger sequencing.

### Growth curve

2.5 µL of overnight cultures were added to 200µL BHIS that had previously been incubated in anaerobic condition overnight to remove dissolved oxygen, without or with 0.5% bile salts in Greiner flat-bottom 96-well plates. A plastic adhesive film (adhesive sealing sheet, Thermo Scientific, AB0558) was added on top of the plate inside the anaerobic station, and the plates were then incubated in a TECAN Infinite M200 Pro spectrophotometer for 24 hours at 37°C. OD600 was measured every 30 minutes, after a 900-second orbital shaking of 2 mm amplitude.

### Determination of minimal inhibitory concentration for vancomycin

*B. thetaiotaomicron* vancomycin E-test was performed on *Brucella* agar supplemented with hemin (5µg/mL), vitamin K1 (1µg/mL) and lysed horse blood (5%v/v). To obtain a bacterial lawn, the agar dishes were covered with a soft-agar lawn composed of *Brucella* agar 0.4% supplemented with hemin (5µg/mL), vitamin K1 (1µg/mL) and lysed horse blood (5%v/v) and inoculated with overnight culture diluted to OD600 = 0.1 (final concentration 10^6^ CFU/mL) in technical triplicates. The vancomycin E-test, purchased from bioMérieux, was placed on the agar plate and incubate at 35°C in anaerobic conditions for 48h. Minimum inhibitory concentrations (MICs) are defined by the lowest concentration of an antimicrobial that will inhibit the visible growth of a microorganism after incubation and were read directly on the graduated scale at the intersection between the inhibition ellipse and the strip (in μg/mL).

### CFU count

The colony-forming unit (CFU) count was performed by a 10 times serial dilution in 200µL of BHIS before and after the PAßN and/or bile salts exposure. A 15µL drop of each dilution for each condition was placed on BHIS plates and incubated at 37°C in anaerobic chamber for 48h before CFU count.

### Hoechst H33342 bisbenzimide accumulation assay

Strains were cultured overnight at 37°C and used to inoculate fresh medium that was incubated for a further 5 h at 37°C. Bacterial cells were collected by centrifugation at 4000 × *g* and resuspended in PBS (1 mL), as previously described (25). The optical density of all suspensions was adjusted to 0.1 at 600 nm and aliquots (180µL) were transferred to Greiner flat-bottom 96- well plates. Eight technical replicates of each strain were analyzed in each column. The plate was transferred to a TECAN Infinite M200 Pro spectrophotometer, incubated at 37°C and H33342 (25 mM) was added (20 µL) to each well to give a final concentration of 2.5 mM. Fluorescence was read from the top of the wells using excitation and emission filters of 346 and 460 nm, respectively, with 5 flashes/well; readings were taken for 30 cycles with a 75 s delay between cycles, and a gain multiplier of 1460. Raw fluorescence values were analyzed using Excel (Microsoft) that included calculation of mean values for each column and subtraction of appropriate control blanks. Three independent experiments were performed.

### 96-well crystal violet biofilm formation assay

Overnight cultures were diluted to OD600 = 0.05 in 150 µL BHIS without or with supplement and inoculated in technical duplicates in polystyrene Greiner round-bottom 96-well plates. The wells at the border of the plates were filled with 200 µL of water to prevent evaporation. Incubation was done at 37°C in anaerobic conditions for 48h. The supernatant was removed by careful pipetting and the biofilms were fixed using 150 µL of Bouin’s solution (picric acid 0.9%, formaldehyde 9% and acetic acid 5%, HT10132, Sigma-Aldrich) for 10min. Then the wells were washed once with water by immersion and flicking, and the biofilm was stained with 175 µL of 1% crystal violet (V5265, Sigma-Aldrich) for 10 minutes. Crystal violet solution was removed by flicking and biofilms were washed twice with water. Stained biofilms were dried then resuspended in 1:4 acetone: ethanol mix and absorbance at 595 nm was measured using TECAN infinite M200 PRO plate reader.

### Extracellular matrix extraction and quantification

Matrix components were extracted based on previously described methods (46). 2mL of 48h biofilms cells grown in presence of bile were harvested and centrifugated at 5000 × *g* for 10 min. The pellet was washed twice with NaCl 0.85% and weighed, then resuspended in an extraction buffer (Tris-HCl pH 8.0; 1.5 M NaCl) at a 1:10 mass-volume ratio and incubated at 25 °C for 30 min with agitation. Then, cells were removed by centrifugation at 15,000 × *g* and 25 °C for 10 min and the supernatant containing the extracted ECMs were stored at -20°C until use. The amount of RNA, DNA and proteins in the ECM were measured using a Qubit 3.0 Fluorometer (Thermo Fisher Scientific) according to the manufacturer’s instructions. The concentration of polysaccharides was quantified by adding to a volume of ECM, in clean and acid washed glass tubes, a 1:1 volume ratio of 5% phenol then 1:5 volume ration of 93% sulfuric acid. After incubation at room temperature for 10 min, 100µL was transferred in polystyrene Greiner flat-bottom 96-well plates prior to OD490 measurement using a TECAN Infinite M200 Pro spectrophotometer. The concentration was then calculated from a standard curve. The eDNA profil were performed adding 1µL of ECM samples in an Agilent DNA chip and analysed using the Agilent 2100 Bioanalyzer system.

### Magnesium quantification assay

Using the Magnesium Assay kit (Sigma-Aldrich, MAK026), the magnesium concentration was determined by a coupled enzyme assay that takes advantage of the specific requirement of glycerol kinase for Mg^2+^, resulting in a colorimetric (450 nm) product proportional to the magnesium present. This assay exhibits no detectable interference with Fe^2+^, Cu^2+^, Ni^2+^, Zn^2+^, Co^2+^, Ca^2+^, and Mn^2+^. Briefly, 25µL of samples is added to 25µL of water, into duplicate in a 96-wells flat bottom plate. In parallel, standard wells containing 0 (blank), 3, 6, 9, 12 and 15 nmole/well standards of magnesium were prepared. Then, 50 µL of the Reaction Mix was added to wells containing samples and standards. The absorbance at 450 nm (A450) was measured every 5 minutes using TECAN infinite M200 PRO plate reader in kinetic mode for 30 minutes at 37°C. The concentration was then calculated from a standard curve.

### Nuclease activity of the supernatant

48h cultures were centrifugated 6.5 min at 6000 × *g*. 50 µl of supernatant was mixed with *B. thetaiotaomicron* VPI-5482 genomic DNA (800 ng/mL final concentration). 10 µL was migrated on a 1% agarose gel and colored with ethidium bromide. The remaining 40 µL was incubated at 37°C overnight for 24h and then 10 µL was used to run a 1 % agarose gel and colored with ethidium bromide.

### Confocal laser scanning microscopy

Biofilms were grown in 96-well plates (μclear, Greiner Bio-One). A total of 150 μL of BHIS, supplemented with 0.5% bile extract, Dnase I (Thermo Scientific, VF304452) 98 U/mL, or Rnase1 (Thermo Scientific, EN0601) 0.06 U/mL when required, was added to each well and the plates were incubated at 37 °C, in static condition 48 h under anaerobic conditions. The unwashed biofilms were then directly stained in red with 20 μM of SYTO61 (Life Technologies; cell permeant nucleic acid dye to contrast all the bacteria) and in green with 0.4 μM of TOTO-1 (Thermo Scientific; cell impermeant DNA dye to contrast eDNA). After 15 min of incubation, Z stacks of horizontal plane images were acquired in 1-μm steps using CLSM (Leica TCS SP8, INRAE MIMA2 microscopy platform) with a water 63× immersion lens (numerical aperture [NA] = 1.2). Two stacks of images were acquired randomly on three independent samples at 800 Hz. Fluorophores were excited and emissions were captured as prescribed by the manufacturer. Simulated 3D fluorescence projections were generated using IMARIS 9.3 soft- ware (Bitplane). Biofilm biovolumes (µm3) extracted from CLSM images were analyzed with BiofilmQ (47).

### Transmission electron microscopy (TEM)

For transmission electron microscopy, biofilms were grown in Falcon® clear PET membrane insert for 12-well plate (ref. 353180) in 12-well plates (Greiner Bio-One). 1mL of BHIS, supplemented with 0.5% bile extract, 25µg/mL PAßN or 35mM magnesium when required, was added to each well and the plates were incubated at 37 °C, in static condition 48h under anaerobic conditions. The unwashed biofilms were then directly fixed in a 0.07 M cacodylate buffer containing 1.3% glutaraldehyde and 0.05% ruthenium red. Samples were then washed in cacodylate-buffer and post-fixed by incubation with 1% osmium tetroxide for 1 h. Samples were then fully dehydrated in a graded series of ethanol solutions and embedded in Epon resin, which was allowed to polymerize from 37°C to 60°C. Ultra-thin sections of these blocks were obtained with an ultramicrotome. Sections were stained with 5% uranyl acetate 5% lead citrate and observations were made with a transmission electron microscope (JEOL 1011, Tokyo, Japan).

### RNAseq analysis

Overnight cultures were mixed with RNAprotect (Qiagen) in the anaerobic chamber to prevent RNA degradation and bacteria were lysed using QIAGEN Proteinase K and TE buffer containing lysozyme. Total RNA was extracted using the Direct Zol kit (Zymo Cat. R2050) according to the manufacturer’s instructions and treated with DNase I from the same kit. RNA concentration, quality, and integrity from 4 independent replicates was checked using RNA6000 Nano chips and the Agilent 2100 Bioanalyzer system. Sequencing was performed by the Biomics platform at the Institut Pasteur. Ribosomal RNA depletion was performed using the Bacteria RiboZero kit (Illumina). From rRNA-depleted RNA, directional libraries were prepared using the TruSeq Stranded mRNA Sample preparation kit following the manufacturer’s instructions (Illumina). Libraries were checked for quality on Bioanalyzer DNA 1000 chips (Agilent). Quantification was performed with the fluorescent-based quantitation Qubit dsDNA HS Assay Kit (Thermo Fisher Scientific). Sequencing was performed as a Single Read run for 75 bp sequences on a NextSeq500 Illumina sequencer. The multiplexing level was 16 samples per lane. The RNA-seq analysis was performed with Sequana (48). In particular, we used the RNA-seq pipeline (v0.15.1, https://github.com/sequana/rnaseq) built on top of Snakemake v6.7.0 (49). The pipeline trimmed reads from adapters and low quality bases using fastp software v0.20.1 (50), then mapped the remaining reads to the *bacteroides thetaiotaomicron* genome (accession NC_004663.1) with bowtie2 v2.4.2 (51). Mapped reads were then quantified using featureCounts (52), and differential analysis performed with DESeq2 v1.30.0 (53) with default parameters. Results are partially presented in supplementary table S2 and presented in full in supplementary table S3.

## ACKNOWLEDGEMENTS

We thank Nathalie Béchon, Sol Vendrell-Fernandez, Christiane Forestier, Muriel Masi, Christophe Beloin and Yutaka Yoshii for critical reading of the manuscript. This work was supported by Institut Pasteur and grants by the ANR DifBiolRel (grant n° ANR 20CE15002201), the French government’s Investissement d’Avenir Program, Laboratoire d’Excellence “Integrative Biology of Emerging Infectious Diseases” (grant n°ANR-10-LABX- 62-IBEID), the *Fondation pour la Recherche Médicale* (grant no. DEQ20180339185). A.A. Lopes was supported by a “Poste d’accueil AP-HP” 2019-2022. The RNA sequencing and analysis was performed by the Biomics Platform, C2RT, Institut Pasteur, Paris, France, supported by France Génomique (ANR-10-INBS-09-09) and IBISA. Biofilm confocal imaging was performed at the INRAE MIMA2 imaging platform.

## AUTHORS CONTRIBUTIONS

A-A.L. and J.-M.G. designed the experiments. A-A.L., J.D., S.G. performed the experiments. A-A.L., T.C., J.D., R.B. and J.-M.G. analyzed the data. Y.Y., A-A.L. and J.-M.G. wrote the paper with significant contribution from all authors.

