## Supporting Figures S1 to S5 and Tables S1, S2 for "Bile-induced biofilm formation in *Bacteroides thetaiotaomicron* requires magnesium efflux by an RND pump"

Supporting Tables S1, S2 (supporting table S3 is provided as an excel spreadsheet uploaded separately as a dataset)

### SUPPLEMENTARY TABLES

**Supplementary Table S1: PAßN does not permeabilize the outer membrane.** Vancomycin susceptibility testing was performed using E-test on VPI-5482 without and with 25 and 50 µg/mL of phenylalanine-arginine ß-naphthylamide (PAßN). Vancomycin is a large antibiotic that is normally incapable of crossing the outer membrane of Gram-negative bacteria without permeabilization of this membrane. The Minimal inhibitory Concentration (MIC) was confirmed by 3 independent experiments for each condition.

| | PAßN (µg/mL) | | |
|---|---|---|---|
| | 0 | 25 | 50 |
| MIC to vancomycin (µg/mL) | >256 | >256 | >256 |

**Supplementary Table S2: RNA-sequencing results comparing gene expression for the 21 *B. thetaiotaomicron* RND-type efflux operons with and without 0.5% bile salts.**

| RND-type efflux pump operons | Genes | Compared log2 fold expression change with and without bile | Adjusted p-value |
|---|---|---|---|
| **BT4693-4695** | BT4693 | -0.219 | 0.71 |
| | **BT4694** | -0.124 | 0.73 |
| | BT4695 | -0.050 | 0.92 |
| **BT3968-3969** | BT3968 | -0.887 | 0.24 |
| | **BT3969** | 0.257 | 0.77 |
| **BT3337-3339** * | BT3337 | 2.615 | <0.001 |
| | **BT3338** | 2.847 | $8.105e^{-11}$ |
| | BT3339 | 2.301 | $6.894e^{-22}$ |
| **BT2940-2942** | BT2940 | -0.164 | 0.88 |
| | **BT2941** | -0.350 | 0.63 |
| | BT2942 | -0.151 | 0.84 |
| **BT2835** * | **BT2835** | 1.716 | 0.0005 |
| **BT2793-2795** * | BT2793 | 7.517 | $7.367e^{-49}$ |
| | **BT2794** | 7.550 | $1.187e^{-68}$ |
| | BT2795 | 7.644 | $2.489e^{-98}$ |
| **BT2686-2688** * | **BT2686** | 3.546 | $1.082e^{-34}$ |
| | BT2687 | 3.270 | $9.385e^{-28}$ |
| | BT2688 | 2.851 | $7.322e^{-79}$ |
| **BT2251-2253** | BT2251 | -0.707 | 0.14 |
| | **BT2252** | 0.261 | 0.42 |
| | BT2253 | -0.119 | 0.46 |
| **BT2117-2119** * | BT2117 | 2.575 | $1.40e^{-6}$ |
| | **BT2118** | 2.595 | 0.00040 |
| | BT2119 | 1.809 | 0.0096 |
| **BT2038-2040** | BT2038 | 0.436 | 0.28 |
| | **BT2039** | -0.023 | 0.97 |
| | BT2040 | -1.074 | 0.04 |
| **BT1965-1967** * | BT1965 | 1.965 | 0.009 |
| | **BT1966** | 2.637 | 0.00005 |
| | BT1967 | 3.142 | 0.021 |
| **BT1693-1695** | BT1693 | -0.930 | 0.17 |
| | **BT1694** | -0.258 | 0.79 |
| | BT1695 | -0.771 | 0.16 |
| **BT1465-1468** | BT1465 | 0.019 | 0.96 |
| | BT1466 | -0.286 | 0.61 |
| | **BT1467** | -1.822 | 0.02 |
| | BT1468 | -1.398 | 0.30 |
| **BT1267-1269** | BT1267 | -0.341 | 0.67 |
| | **BT1268** | 0.010 | 0.99 |
| | BT1269 | -0.999 | 0.58 |
| **BT0884-0886** | BT0884 | -0.532 | 0.55 |
| | BT0885 | -0.265 | 0.71 |
| | **BT0886** | -0.419 | 0.47 |
| **BT0678-0680** | BT0678 | -0.166 | 0.87 |
| | BT0679 | 0.235 | 0.82 |
| | **BT0680** | -0.044 | 0.97 |
| **BT0669-0672** | BT0669 | -1.017 | 0.05 |
| | **BT0670** | 0.070 | 0.91 |
| | BT0671 | 0.939 | 0.39 |
| | **BT0672** | -0.254 | 0.85 |
| **BT0304-0306** | BT0304 | -1.267 | 0.0003 |
| | **BT0305** | -0.2794 | 0.69 |
| | BT0306 | -0.308 | 0.63 |
| **BT0297-0300** * | BT0297 | 2.139 | 0.06 |
| | BT0298 | 2.799 | 0.009 |
| | **BT0299** | 1.202 | 0.109 |
| | **BT0300** | 2.0378 | 0.065 |

* *B. thetaiotaomicron* VPI-5482 operon deleted in this study.
Gene name **in bold** in the gene column correspond to the RND-type efflux inner pump gene.

**Supporting Table S3** (provided as an excel spreadsheet dataset). RNAseq analysis: table of A. all genes, B. upregulated, C. downregulated genes in presence of 0.5% bile, D. COG functional categories enrichment.

### SUPPLEMENTARY FIGURES

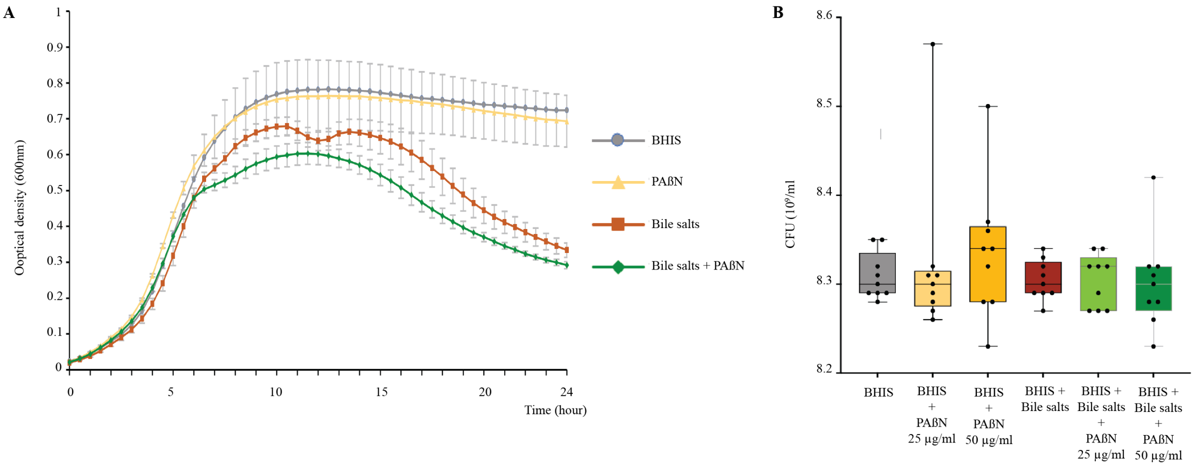


**Figure S1: PAßN does not affect growth nor viability. A.** 24 hours growth curve of VPI-5482 in BHIS without or with 25µg/mL of PAßN and/or 0.5% bile salts (BS) in 96-well plate. Mean of 6 biological replicates, error bars represent standard deviation to the mean (SEM). **B.** Quantification of cells in overnight cultures of VPI-5482 grown in BHIS with or without phenylalanine-arginine ß-naphthylamide (PAßN) at 25 or 50 µg/mL and/or 0.5% bile salts (BS). Min-max boxplot of 9 biological replicates for each condition. Statistics correspond to an unpaired, nonparametric Mann–Whitney *U* test. No significant difference was detected (p-value > 0.05).

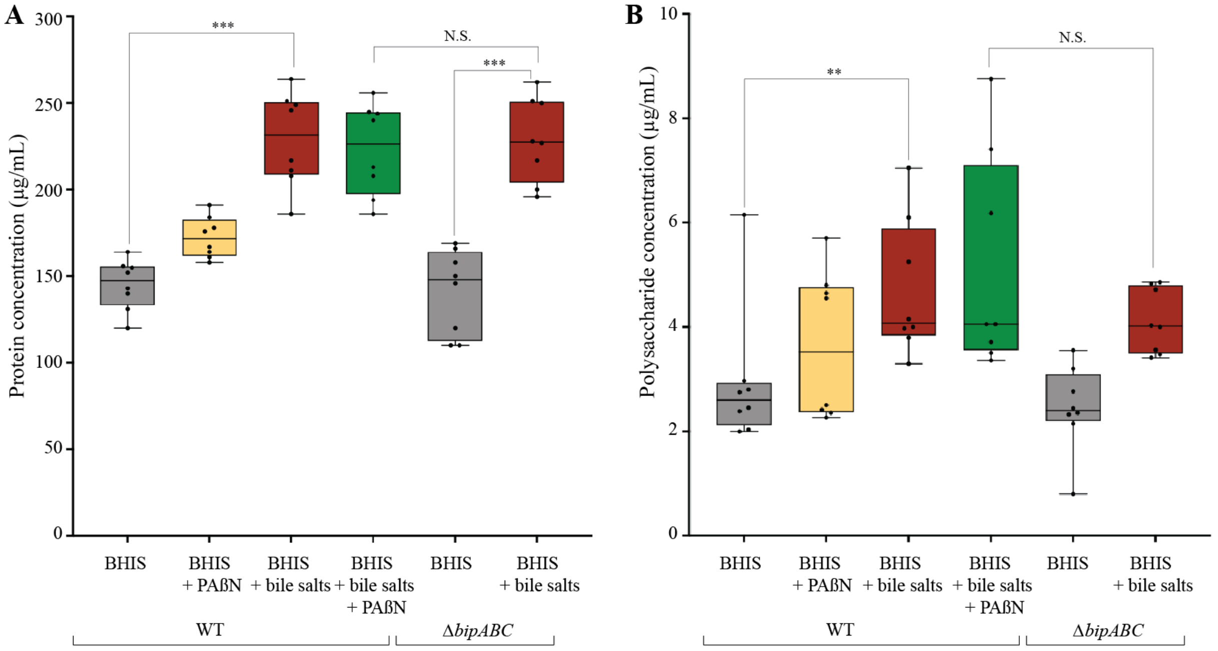


**Supplementary Figure S2: Impact of impaired efflux on extracellular matrix protein and polysaccharide composition.** Concentration of proteins (**A**) and polysaccharides (**B**) in the ECM of *B. thetaiotaomicron* VPI-5482 WT or its corresponding Δ*bipABC* mutant without and with 0.5% bile salts (BS) or 25 µg/mL of PAßN. Min-max boxplot of 8 biological replicates for each condition. ** p-value<0.005, *** p-value<0.0005. N.S.: non-significant. Statistics correspond to an unpaired, nonparametric Mann–Whitney *U* test.

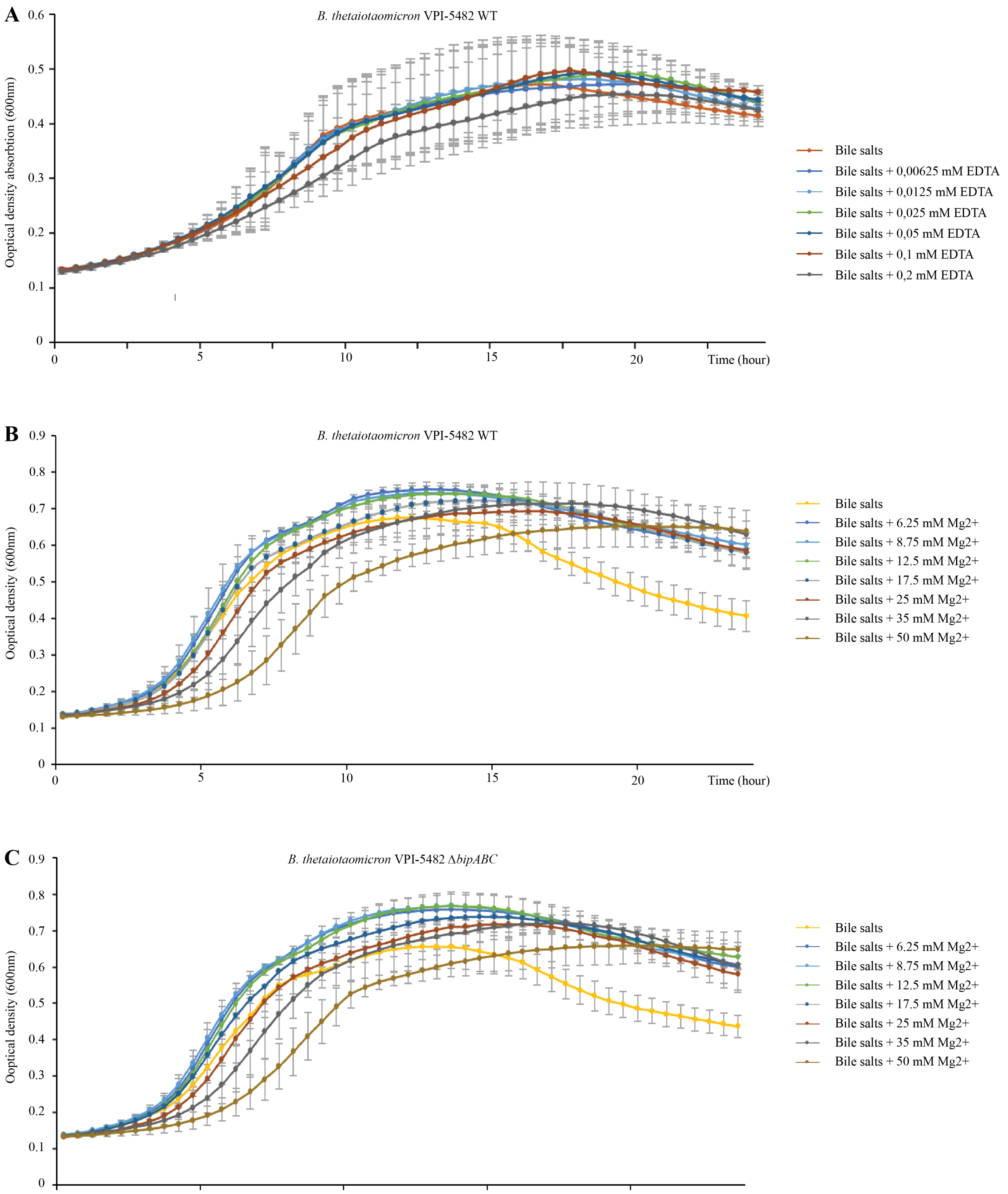


**Supplementary Figure S3: Non-toxicity of EDTA and Magnesium on the strains used. A.** 24 hours growth curve of *B. thetaiotaomicron* VPI-5482 in BHIS with 0.5% bile salts and different concentrations of EDTA from 0 to 0.2 mM in 96-well plate. **B and C.** 24 hours growth curve in BHIS with 0.5% bile salts and different concentrations of Magnesium from 0 to 50 mM in 96-well plate in *B. thetaiotaomicron* VPI-5482 WT (**B**) and Δ*bipABC* (**C**). Mean of 4 biological replicates, error bars represent standard deviation to the mean (SEM).

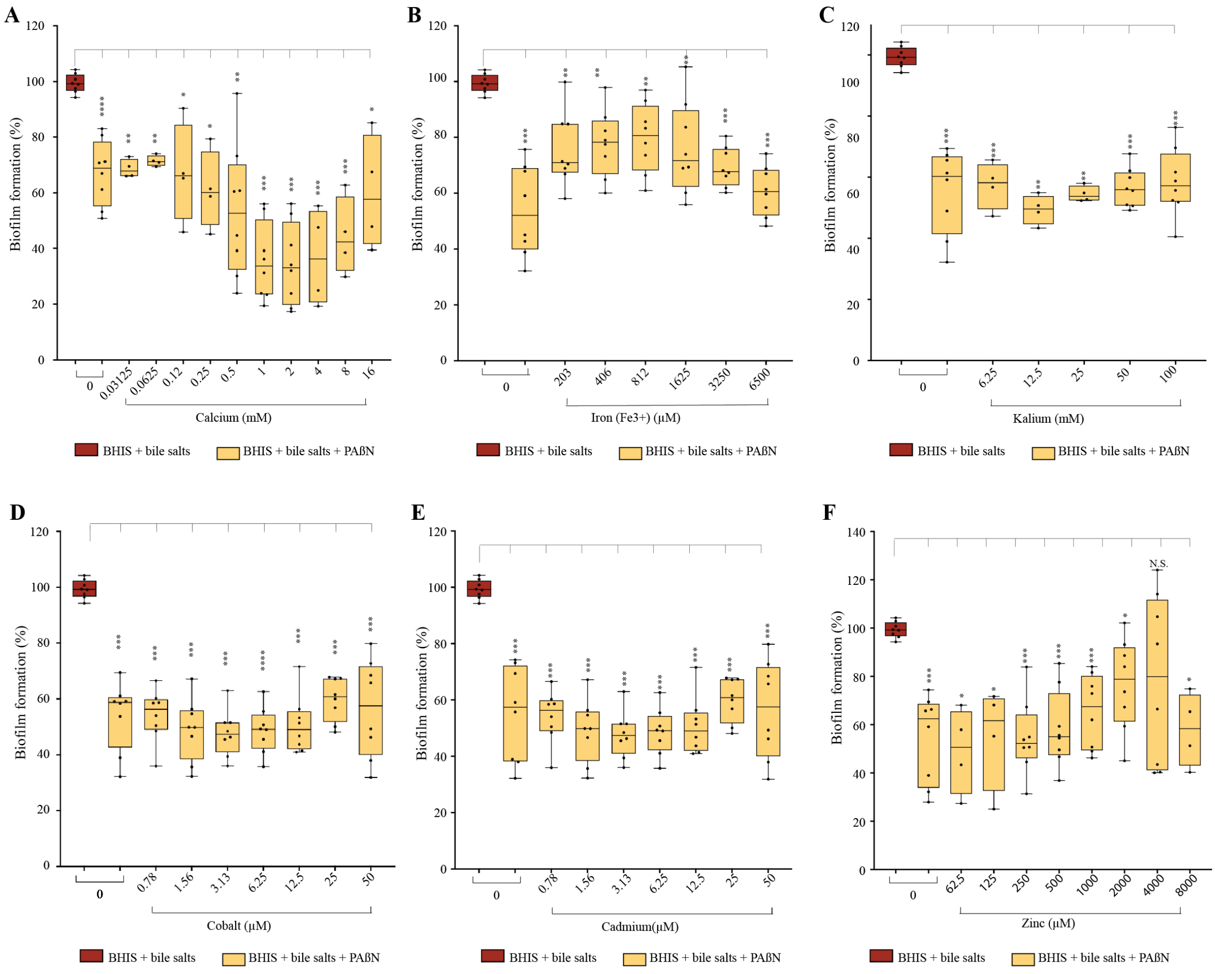


**Supplementary figure S4: Addition of cations other than magnesium does not restore biofilm formation of VPI-5482 with PAßN.** 96-well plate crystal violet biofilm assay after 48h growth in BHIS of VPI-5482 in the presence of 0.5% bile salts (BS), without or with 25µg/mL of PAßN and different non-toxic concentrations of calcium (**A**), iron (**B**), kalium (**C**), cobalt (**D**), cadmium (**E**) and zinc (**F**). Mean of WT in BHIS with 0.5% BS was adjusted to 100%. Min-max boxplot of 8 biological replicates for each condition. * p-value<0.05; ** p-value<0.005; *** p-value<0.0005; **** p-value<0.00005 compared to WT in BHIS with BS. N.S.: non-significant. Statistics correspond to an unpaired, nonparametric Mann–Whitney *U* test.

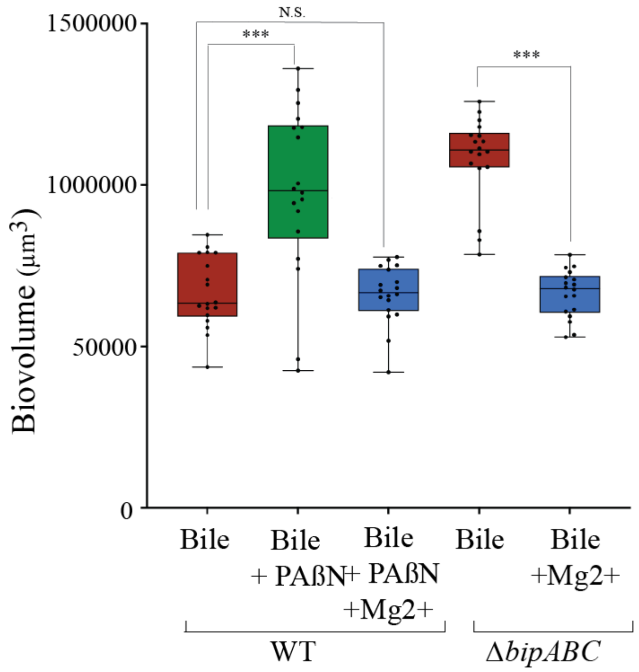


**Supplementary figure S5: Increased biovolume upon addition of PAßN and deletion of the BipABC RND pump is complemented by supplementation with magnesium**. Biofilm biovolumes (µm$^3$) extracted from CLSM images were analyzed with BiofilmQ. Each value is a mean of 18 values obtained from 3 independent experiments. Statistics correspond to an unpaired, nonparametric Mann–Whitney *U* test.
